# Global Marine Cold Seep Metagenomes Reveal Diversity of Taxonomy, Metabolic Function, and Natural Products

**DOI:** 10.1101/2023.04.06.535842

**Authors:** Tao Yu, Yingfeng Luo, Xinyu Tan, Dahe Zhao, Xiaochun Bi, Chenji Li, Yanning Zheng, Hua Xiang, Songnian Hu

**Author notes:** Corresponding authors. (Hu S), (Xiang H). Equal contribution (Tao Yu, Yingfeng Luo).

## Abstract

Cold seeps in the deep sea are closely linked to energy exploration as well as global climate change. The alkane-dominated chemical energy-driven model makes cold seeps an oasis of deep-sea life, showcasing an unparalleled reservoir of microbial genetic diversity. By analyzing 113 metagenomes collected from 14 global sites across 5 cold seep types, we present a comprehensive Cold Seep Microbiomic Database (CSMD) to archive the genomic and functional diversity of cold seep microbiome. The CSMD includes over 49 million non-redundant genes and 3175 metagenome-assembled genomes (MAGs), which represent 1897 species spanning 106 phyla. In addition, beta diversity analysis indicates that both sampling site and cold seep type have substantial impact on the prokaryotic microbiome community composition. Heterotrophic and anaerobic metabolisms are prevalent in microbial communities, accompanied by considerable mixotrophs and facultative anaerobes, indicating the versatile metabolic potential in cold seeps. Furthermore, secondary metabolic gene cluster analysis indicates that at least 98.81% of the sequences encode potentially novel natural products. These natural products are dominated by ribosomal processing peptides, which are widely distributed in archaea and bacteria. Overall, the CSMD represents a valuable resource which would enhance the understanding and utilization of global cold seep microbiomes.

## Introduction

Marine cold seep is a special chemoenergetic trophic ecosystem driven by gaseous and liquid hydrocarbon from deep geologic sources [1,2]. Despite such extreme environment of low oxygen and temperature, high pressure, and absence of light [3], the anaerobic methanotrophic archaea (ANME) and sulfate reducing bacteria (SRB), which are dominated by the utilization of methane and other alkanes [4–7]. Methane-dominated short-chain alkanes released from cold seep may enter the atmosphere and thus affect the global climate, accompanied by natural leakage processes and human mining activities [8]. In addition, mining activities may negatively affect the biodiversity at regional and global scales by disrupting the original microbial communities of cold seep [9]. Therefore, understanding the microbiome composition associated with cold seeps is critical for addressing the global energy crisis and climate change, as well as for utilizing the microbial resources of cold seep.

In recent years, with advances in high-throughput sequencing technologies and computational methods, several comprehensive metagenomic databases have been constructed, including glacier [10], marine [11], human [12], and Earth microbiomes [13]. These studies have promoted substantial understanding of microbial community composition, and the metabolic properties of microbiome in specific habitats. Although, for cold seep, there are studies related to microbial community composition [2,14,15], carbon cycling [5,7,16–18], and nitrogen cycling [19,20], a comprehensive and complete database integrating all known global cold seep samples is still lacking. This inevitably limits the systematic understanding of cold seep microbiome.

Furthermore, because cold seeps possess a rich species diversity and the vast majority of species are uncultured, they may harbor tremendous phylogenetic, metabolic, and functional diversity. Natural products produced by diverse secondary metabolite biosynthetic gene clusters (BGCs) mainly include non-ribosomal peptide synthases (NRPSs), polyketide synthases and their derivatives (PKSI and PKS other), PKS-NRPS hybrids, ribosomal processing peptides (RiPPs), and terpenes [21]. It has been widely demonstrated to have substantial value in medicine, agriculture, and biotechnology [22,23]. For example, from 1981 to 2019, 36.3% of new drugs approved by the US Food and Drug Administration were natural products or their derivatives [22]. Numerous studies have shown that a large number of uncultured microbiome encoding BGCs in land [13], marine [11,13] and glacier [10]. However, the biosynthetic potential of cold seep microbiome remains largely unexplored.

Currently, scattered and non-uniform metagenomic studies limit the understanding of the microbial diversity of the global cold seep ecosystem. Accordingly, we performed an integrative analysis of 113 metagenomes from 14 global sites covering five cold seep types. Here, we present the prokaryote-focused Cold Seep Microbiomic Database (CSMD). The catalog includes 1897 potential species-level prokayotic genomes derived from 3175 metagenome-assembled genomes (MAGs), 27 M contigs, and over 49 M non-redundant genes, thus facilitating the exploration of global cold seep microbial composition and metabolic diversity, as well as the assessment of natural product synthetic potential in particular.

## Results

### Construction of CSMD

We obtained a total of 113 metagenomic samples from 14 cold seep sites globally, comprising 101 publicly available samples and 12 samples we collected. These sites encompassed five distinct types of seepage: methane seep, oil and gas seep, gas hydrate, asphalt volcano, and mud volcano (**Figure 1A**; Table S1). Metagenomic assembly and binning produced 4335 MAGs, which were combined and de-redundant with publicly available 1688 MAGs to finally obtain 3175 MAGs (Figure 1B and Table S2). All of them meet the medium and above quality level of the Minimum Information about a Metagenome-Assembled Genome (MIMAG) criteria (completeness ≥ 50%, contamination < 10%) [24] with a mean completeness of 71.24% (± 13.45%) and a mean contamination rate of 3.77% (± 2.78%) (Figure 1C). The microbial genomes of cold seep harbor diverse genome size (0.50 Mb to 9.26 Mb) and GC content (23.14 % to 72.66%). In addition, 49.87 % and 99.94 % of total genomes were identified at least one ribosomal RNA (rRNA) and transfer RNA (tRNA) gene fragments (Table S2).

**Figure 1.**
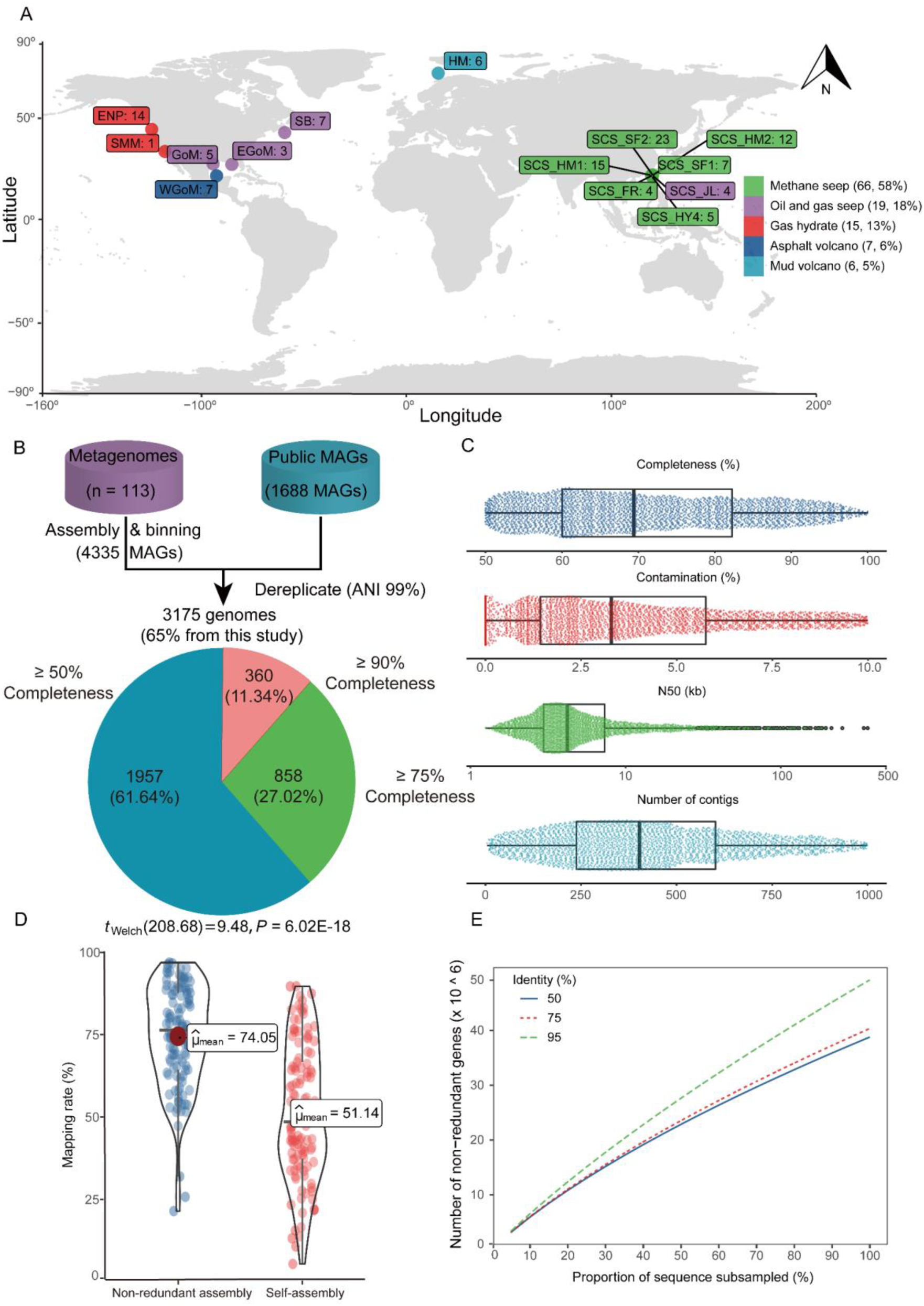
Construction of global CSMD. **A.** Geographic distribution of cold seep metagenomes. **B**. A total of 3175 cold seep genomes were recovered from metagenomes and public MAGs. **C**. Distribution of quality metrics across genomes (*n* = 3175), showing the minimum value, first quartile, median, third quartile and maximum value. **D.** Distribution of sample reads mapping rate against 56 Gb non-redundant assembly and self-assembly. Welch’s t-test was performed for two groups. E. Gene diversity analysis based on 50%, 75%, and 95% nucleotide identity. CSMD, Cold Seep Microbiomic Database; MAGs, metagenome-assembled genomes.

Additionally, 113 assembled metagenomes were merged and de-redundant, resulting in a 56 Gb non-redundant contigs of the cold seep microbiome after removing eukaryotic contigs annotated by CAT [25]. A total of 27,599,955 contigs with a mean length of 2.03 kb and a N50 size of 2.08 kb were comprised in this catalog. Among these 56 Gb non-redundant contigs, 73.88%, 13.64% and 0.24% were taxonomically annotated as bacteria, archaea and viruses respectively (**Figure 2A–C**) via CAT annotation [25] based on NCBI non-redundant protein database. Proteobacteria, Chloroflexi, Bacteroidetes, Planctomycetes, and Acidobacteria were the top 5 most abundant phyla among bacteria, accounting for 39.74% of total contigs (Figure 2A). Euryarchaeota, Candidatus Lokiarchaeota, Candidatus Bathyarchaeota, Candidatus Thorarchaeota, and Candidatus Heimdallarchaeota were the top five most abundant archaea, accounting for 5.49% of total contigs, while Uroviricota, Nucleocytoviricota, Cressdnaviricota, Preplasmiviricota, and Phixviricota were the top five most abundant viruses, accounting for 0.13% of total contigs (Figure 2B and C).

**Figure 2.**
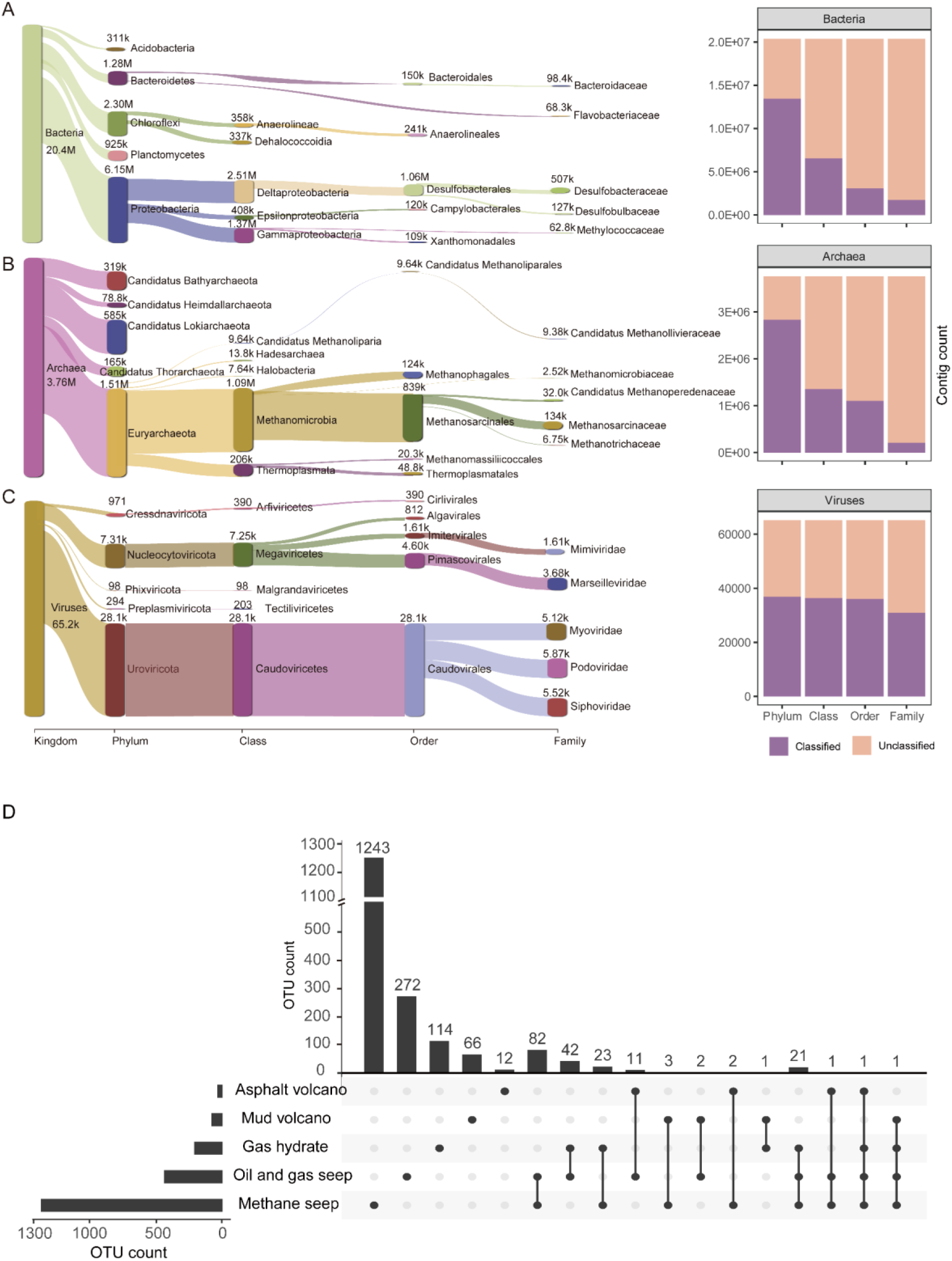
Taxonomic annotation of 56 Gb non-redundant contigs and 1897 OTUs distributed across cold seep types. Sankey plot based on assigned taxonomy showing the dominant (left) and novel (right) populations at different phylogenetic levels, with the top 5 taxa shown for each level. Numbers indicate the number of contigs for the lineage. **A.** Bacteria. **B.** Archaea. **C.** Viruses. **D.** OTUs intersections across sample groups. An UpsetPlot illustrates OTUs intersections among cold seep types. OTUs, operational taxonomic units.

The self-mapping analysis showed that a range of 4.92% to 89.23% of reads could be mapped to the respective assembly, with an average mapping rate of 51.14 ± 20.40% (Figure 1D and Table S3). Nevertheless, when using the 56 Gb non-redundant contigs as the reference, the average mapping rate increased to 74.05 ± 15.45% (15% to 96%), representing a 23% improvement in mapping rate on average. Therefore, the catalog could be a fundamental reference to facilitate cold seep metagenomic analysis in the future.

Furthermore, a non-redundant protein-coding gene catalog of 49,223,463 gene clusters representing 71,499,869 full or partial length genes was compiled with an alignment percentage threshold of 80% and a nucleotide identity threshold of 95% [10], with 18.69% gene cluster containing at least two members. The gene representatives of 33.55% cluster were complete based on Prodigal [26] prediction. With increasing depth of sampling, the number of non-redundant genes increased steadily and didn’t reach a plateau even at 50% nucleotide identity threshold (Figure 1E). This implies that the cold seeps harbor a substantial genetic diversity, necessitating further sequencing efforts to comprehensively capture its functional diversity. Swiss-Prot [27], UniRef50 [28] and NR databases were used to annotate the functions of de-replicated gene clusters, 33.28%, 79.15% and 80.26% of genes were hit, respectively. These results suggest that the cold seep microbiome has the potential to encode numerous novel proteins.

### Overview of microbiome composition in cold seeps

By combining an average nucleotide identity (ANI) threshold of 95% with an alignment coverage threshold of 30%, 3175 MAGs were clustered into 1897 operational taxonomic units (OTUs) at species level (Table S2). The 1897 OTUs exhibited low sequence identities with genomes from other environmental bacterial and archaeal genomic databases according to the threshold of 95 % ANI, including Tibetan Glacier Genomes (TGG) (100% novelty) [10], the TaraOcean genomes (100% novelty) [20], the Ocean Microbiomics Database (OMD) (99.21% novelty) [11], Genomes from Earth’s Microbiomes (GEM) (97.79% novelty) [13], and Genome Taxonomy Database (GTDB) (94.41% novelty) [29] (Table S4). Approximately 90% (1707 OTUs, 89.98%) of the OTUs were present in only one cold seep type, 8.75% (166 OTUs) in two types, and only 1.27% (24 OTUs) in three or more types (Figure 2D and Table S5). Similarly, the abundance of OTUs showed a high degree of niche specialization of cold seep (Figure S1 and Table S6). Thus, further investigation of cold seep microbial diversity is necessary.

According to the GTDB (release R06-R202) [30] annotation, the cold seep genomic dataset shows a substantial taxonomic diversity. The 1897 OTUs span across 106 phyla, 173 classes, 308 orders, 433 families and 407 genera (Table S2). In addition, the number of species in 17 under-represented phyla were expanded for 1.25-4 times compared to GTDB R202 [29] (Table S7). For example, uncultured UBP7_A was increased to 4-fold, Krumholzibacteriota was increased to 2.1-fold, and Asgardarchaeota was increased to 1.28-fold (Table S7). Further, 46 classes, 130 orders, 297 families, 960 genera, and 1790 species represent potential novel lineages compared to the GTDB. Chloroflexota (200 OTUs, 10.06%), Proteobacteria (197 OTUs, 9.91%), Desulfobacterota (145 OTUs, 7.29%), Planctomycetota (106 OTUs, 5.33%) and Patescibacteria (106 OTUs, 5.33%) were the five phyla of bacteria that contain a relatively high number of species, while Halobacteriota (75 OTUs, 3.77%), Asgardarchaeota (74 OTUs, 3.72%), Thermoplasmatota (68 OTUs, 3.42%), Thermoproteota (52 OTUs, 2.62%) and Nanoarchaeota (40 OTUs, 2.01%) were the phyla of archaea that contain a relatively high number of species (Table S2). Even with 296 high quality OTUs (completeness > 90%), 10 classes, 21 orders, 38 families, 124 genera and 254 species represented potential novel lineages (Table S2).

The fluid systems of cold seeps are usually classified as mineral-prone systems (*e.g*., methane seep, oil and gas seep, and gas hydrates) with low discharge and mud-prone systems (*e.g*., mud volcano and asphalt volcano) with high discharge, according to the fluid flow regime [1]. Simpson and Shannon diversity of the cold seeps microbiome were significantly higher in mineral-prone systems than in mud-prone seep systems based on OTUs (Figure S2B and Figure S3). Furthermore, we investigated the microbial composition across sampling sites and cold seep types based on the relative abundance of OTUs. In terms of the average relative abundance, Halobacteriota (18.74%), Desulfobacterota (15.2%), Chloroflexota (12.07%), Caldatribacteriota (9.47%), and Proteobacteria (8.48%) represented the most abundant phyla (Figure S2A). Meanwhile, PCoA analysis of microbial communities using Bray-Curtis distance, showed that sampling sites had greater impacts on the distribution of microbiome communities than that in cold seep types at the phylum level of MAGs (Figure S1C) and 16S (Figure S4 and Table S8).

### Versatile metabolic potential of the CSMD

To study the metabolic potential of cold seep microbiome, 1897 OTUs were functionally annotated based on kyoto encyclopedia of genes and genomes (KEGG) database. We first investigated Anaerobic Oxidation of Methane (AOM), a metabolic process which is a primitive driver of the cold seep ecosystem. We found that 30 OTUs contained the marker genes of AOM pathway. Among them, 13 of which had the complete genes involved in the oxidation of methane to CO_2_, 29 had the complete genes from methane to acetate, and 12 had both metabolic steps (**Figure 3** and Table S9). All these 30 OTUs were affiliated to ANME, and 22 of which represented novel genera or species. Additionally, we also found that 1163 OTUs (61.31%; 90 phyla) contained at least one of five pathways for CO_2_ fixation (Figure 3 and Table S9). Among them, the Wood-Ljungdahl pathway (WL pathway) (636 OTUs) was the most prevalent, followed by the Calvin-Benson-Bessham cycle (CBB) (402 OTUs), the 3-hydroxypropionate bi-cycle (3-HP) (170 OTUs), reverse tricarboxylic acid cycle (rTCA) (107 OTUs), 3-hydroxypropionate-4-hydroxybutyric acid cycle (3-HP/4-HB cycle) (240 OTUs) (Figure 3 and Table S9). The top 5 most widely distributed phyla harboring WL pathway were Chloroflexota, Desulfobacterota, Planctomycetota, Thermoplasmatota, and Halobacteriota. The WL pathway in bacteria has been widely discovered, with experimental validation or computational inference in Chloroflexota [31] and Desulfobacterota [32] in the ocean. Compared to other organisms, the WL pathway in archaea is poorly understood [33]. A recent study has shown that Thermoplasmatota have the ability to perform autotrophic growth via the WL pathway [34], and we have also identified 27 OTUs belonging to Thermoplasmatota that possess this pathway. Interestingly, we found that 22 OTUs belonging to Halobacteriota possess key enzymes for the WL pathway, which has not been reported before. Further experiments are required to confirm this *in silico* observation. To explore the heterotrophic potential of the cold seep microbiome, we investigated the genes involved in carbohydrate degradation, and we found that 1887 OTUs may perform heterotrophic metabolism (Figure 3 and Table S9). A novel species from Planctomycetota (SRR13892593_me2_bin.111) possessed the most numerous genes (145 genes) for carbohydrates degradation. Totally, 1163 OTUs (61.31%) belonging to 90 phyla that encoded both the carbohydrate-degrading enzymes and any of the inorganic carbon fixation pathway were considered as potential mixotrophs [35], albeit not rigorously so.

**Figure 3.**
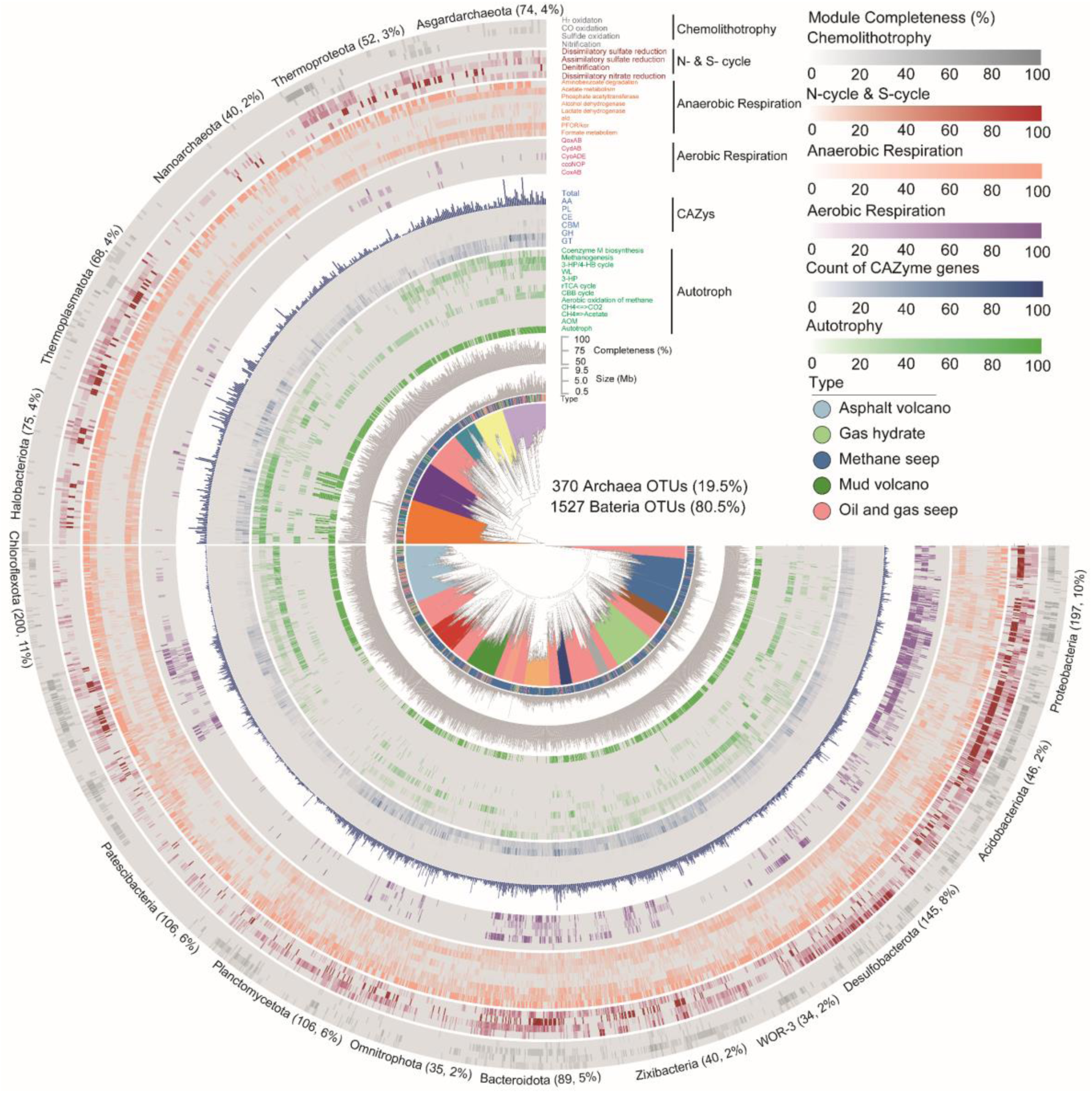
Heat-map illustration of phylogenomic distribution and metabolic profile for 1897 OTUs in the CSMD. The phylogenetic tree was inferred using IQ-TREE from an aligned concatenated set of 120 single copy marker proteins for bacteria, and from a concatenated set of 122 marker proteins for archaea. The key genes of the following pathway are displayed in the diagram: Anaerobic Oxidation of Methane (AOM); Carbohydrate Active Enzyme (CAZy), including glycosidases or glycosyl hydrolases (GH), glycosyltransferases (GT), polysaccharide lyases (PL), carbohydrate esterases (CE), auxiliary activities (AA), and carbohydrate-binding modules (CBM); CO_2_ fixation, including Wood-Ljungdahl pathway (WL), Calvin-Benson-Bessham cycle (CBB), reverse tricarboxylic acid cycle (rTCA), 3-hydroxypropionate bi-cycle (3-HP), and 3-hydroxypropionate/4-hydroxybuty cycle (3-HP/4-HB); anaerobic respiration; aerobic respiration, and chemolithotrophy (refer to Table S9 for details).

Oxygen requirement analysis revealed that all OTUs had at least one anaerobic respiratory pathway. As an illustration, our analysis found the presence of 1296 OTUs with formate metabolism, 582 OTUs with lactate dehydrogenase, 752 OTUs with alcohol dehydrogenase, 1583 OTUs with acetate metabolism, and 1116 OTUs with aminobenzoate degradation (Figure 3 and Table S9). Furthermore, we investigated the potential of the cold seep microbiome to perform aerobic respiration. In total, 736 OTUs (38.79%) were found to contain aerobic respiration genes, such as cytochrome c oxidases (Cox/Cyd/Qox/cco/Cyo) genes (Figure 3 and Table S9). These OTUs are associated with 44 phyla such as Proteobacteria, Asgardarchaeota, Halobacteriota, Chloroflexota, and Nanoarchaeota. All 736 OTUs may perform aerobic respiration using at least one of the anaerobic respiration pathways, indicating potential facultative anaerobic capabilities. These species span 55 phyla, primarily including Proteobacteria (181 OTUs), Chloroflexota (101 OTUs), Bacteroidota (78 OTUs), and Desulfobacterota (54 OTUs). Taking together, cold seep microorganisms was prevalent for anaerobic respiration, and accompanied with substantial genes involved in aerobic respiration.

### Biosynthetic potential of the CSMD

To explore the value of the CSMD, we analyzed the potential of synthesizing natural products. We identified 17,968 putative BGCs with an average length of 7.85 k (± 6.96 k) from the cold seep assemblies using AntiSMASH (version 5.1) [36] (Table S10). To reduce the effect of incomplete and redundant BGCs, these BGCs were clustered into 9390 gene cluster families (GCFs) with an average length of 8.54k (± 7.63k). This was nearly 3.75 times of function-known BGCs within Minimum Information about a Biosynthetic Gene (MIBiG) (https://mibig.secondarymetabolites.org/stats) [37], demonstrating the high diversity of BGCs in the cold seep microbiome. 29.61% of the GCFs had 2 or more members (Table S10). A total of 3112 (33.14%) GCFs containing NRPSs and PKSs were identified from 70 phyla (Table S10 and Figure S5). 3082 (32.82%) GCFs containing RiPPs were identified from 17 phyla, and 845 (8.99%) GCFs containing terpenoids were identified from 10 phyla, with the above total accounting for 75% (Table S10 and Figure S5). This may be due to the fact that among these types, RiPP-like, ranthipeptide, and thiopeptide BGCs may be widely involved in quorum sensing, osmotic stress, and the regulation of cellular metabolism in cold seep microorganisms [11,38]. It is noteworthy that BGC types show substantial variation among different phyla, but their distribution across cold seep types appears to be relatively consistent (**Figure 4A**). This could be attributed to the fact that different phyla commonly carry genes that encode particular natural products. For instance, Chloroflexota and Planctomycetota frequently possess genes that encode genes involved with terpene [10,13], whereas Firmicutes typically harbor genes that encode genes associated with NRPS [11,39].

**Figure 4.**
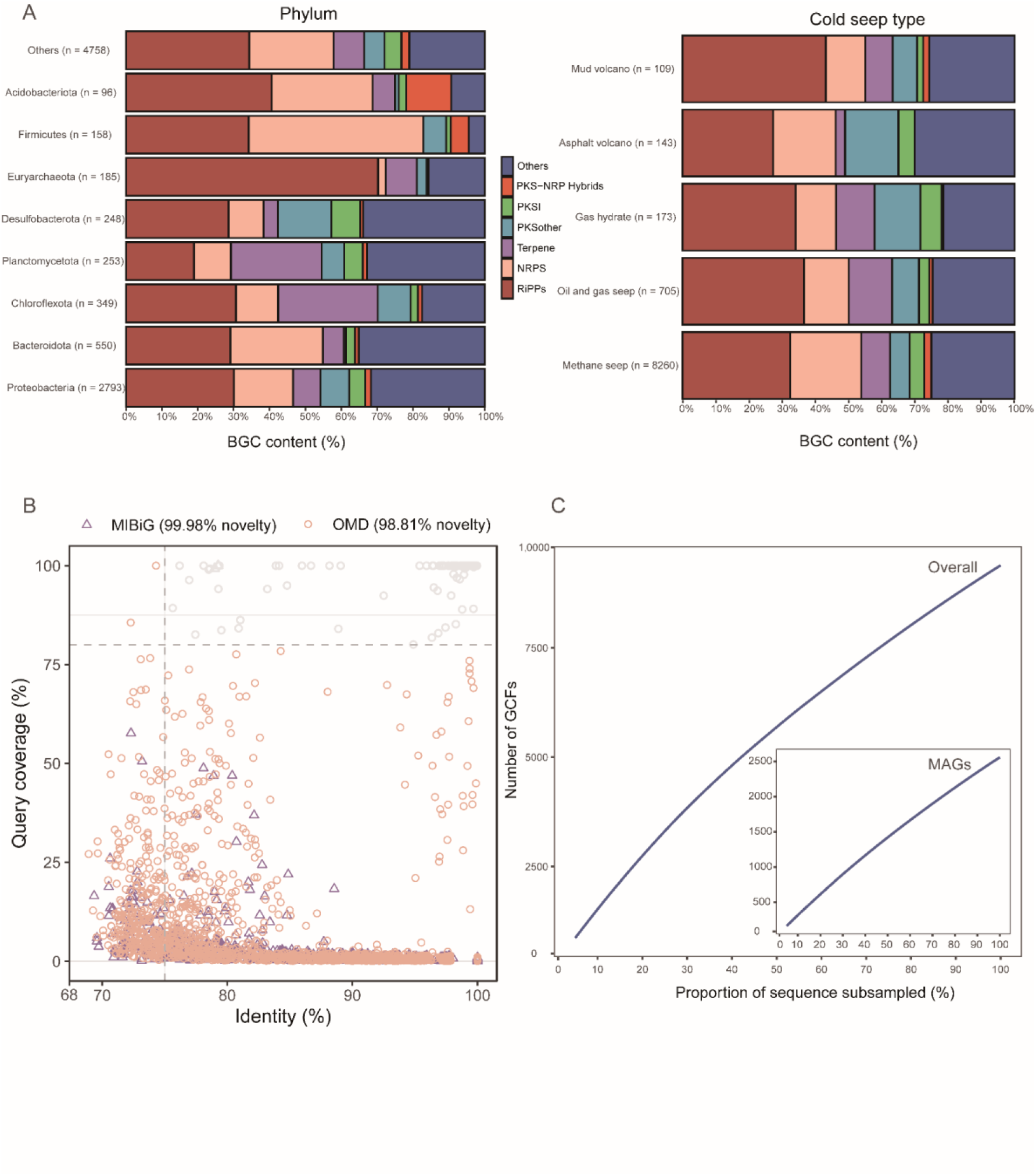
The diversity and novelty of BGCs identified in cold seep microbiomes. **A.** The relative frequency of BGC classes across dominant phyla (left) and cold seep types (right). **B.** Comparing GCFs to experimentally validated MIBiG and computationally predicted (OMD) BGCs uncovers the novelty of GCFs. Only results with BLASTN E-value less than 1E-5 were shown. **C.** Rarefaction curves of GCFs derived from all contigs and MAGs. BGCs, biosynthetic gene clusters; MIBiG, minimum information about a biosynthetic gene; OMD, Ocean Microbiomics Database; GCFs, gene cluster families.

To assess the novelty of the BGCs identified in this study, we compared representative GCFs to MIBiG and OMD. By using 80% query coverage and 75% identity via BLASTN [40], only 2 GCFs were identified in MIBiG and 98.81 % (9278) of the GCFs were considered as novel BGCs compared to OMD (Figure 4B), which may encode novel chemical components. For example, one PKS-NRP hybrids clusters of 84,733 bp comprising 10 core modules was identified from a MAG (SRR13892603_vb_S1C4173) classified as novel genus in family UBA2199 (Figure S5A) showed the most similar (71% Amino Acid Identity (AAI)) to the antibiotic sevadicin biosynthesis gene cluster of *Paenibacillus larvae*. Another RiPPs cluster of 44,319 bp comprising 4 core modules was identified from a MAG (SRR13892601_vb_S1C33830) classified as novel species of Omnitrophota (Figure S5B) showed the most similar (28% AAI) to the antibiotic ranthipeptide of *Streptococcus mutans UA159*. In addition, as sampling BGCs increased, the number of GCF was steadily increased, whether originating from MAGs or contigs (Figure 4C), suggesting that BGCs in cold seep were subject to further exploration, which was in line with the trend of the taxonomic exploration.

### Phylogenetic distribution of BGC-rich clades

To better reveal the relationship between cold seep microbial taxonomy and natural products, we mapped the phylogenetic distribution of BGC-rich clades. For this purpose, 3175 MAGs were placed in GTDB’s standardized bacterial and archaeal phylogenetic trees and overlaid the number of BGCs types (**Figure 5A** and B; Table S11). Totally, 45.92 % (1458) of MAGs contained at least one BGCs with an average length of 9.8 k (± 8.8 k). Overall, bacteria (2.38 ± 1.94) had a higher BGC count per genome than archaea (1.28 ± 0.62) (*P* < 0.001, Mann−Whitney test, Table S11). Furthermore, even after normalizing for genome size, bacteria had a higher BGC count per Mb (1.01 ± 0.7) than archaea (0.78 ± 0.39) (*P* < 0.00001, Mann−Whitney test, Table S10). The results indicate that bacteria have a greater potential for synthesizing natural products than archaea. MAGs with Proteobacteria, Desulfobacterota, Bacteroidota, Chloroflexota, and Planctomycetota are the bacterial phyla with the highest number of BGCs, consistent with the predictions based on all contigs (Figure 4A). In addition, 238 BGCs were detected within Halobacteriota (110 BGCs), Thermoplasmatota (45 BGCs), Asgardarchaeota (36 BGCs), Thermoproteota (34 BGCs), and Nanoarchaeota (13 BGCs), dominant by RiPPS, NRPS and PKS group (Table S11). Overall, the CSMD provides access to novel lineages, including microbial resources for the discovery of novel natural products.

**Figure 5.**
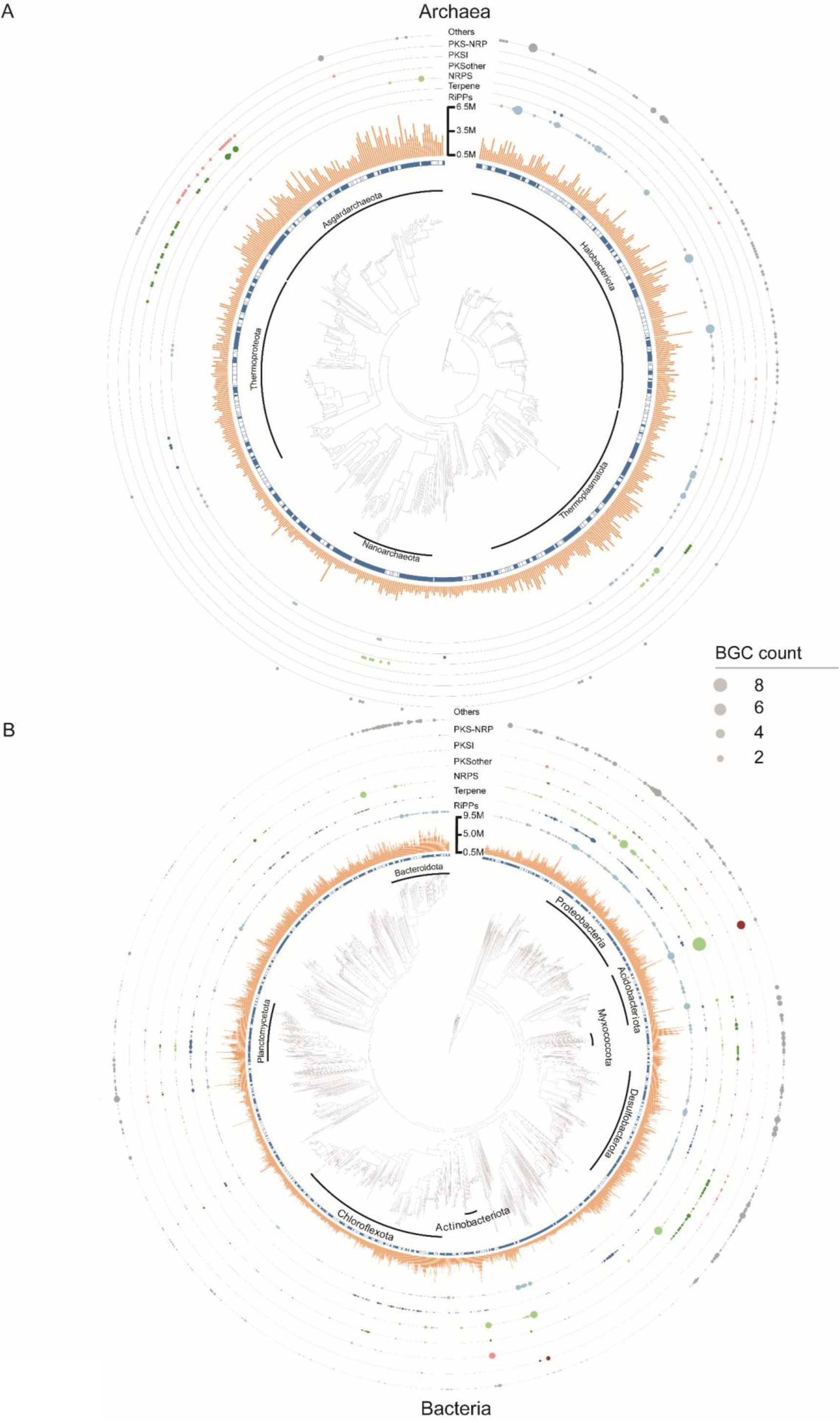
Illustration of BGC-rich lineages in cold seep microbiomes. The solid square in the innermost circle indicates the representative genome of each OTU. The circle size indicates the number of BGCs for each category. **A.** Archaea. **B.** Bacteria.

Afterwards, to investigate the overlap of the natural products among different phyla and cold seep types, we examined the distribution of GCFs within each phylum and cold seep types (**Figure 6A** and B). In most phyla, the majority (73.81% ± 20.35%) of GCFs appeared to be phylum-unique (Figure 6A). Likewise, the shared GCFs are rarely observed among cold seep types, with the majority of GCFs were detected in only one type (Figure 6B). Exceptionally, there are a few shared GCFs between methane seep and oil/gas seep, which are observed in MAGs (Figure 6B) and samples (Figure 6C). This may be due to that these two types have similar environmental factors [1,2].

**Figure 6.**
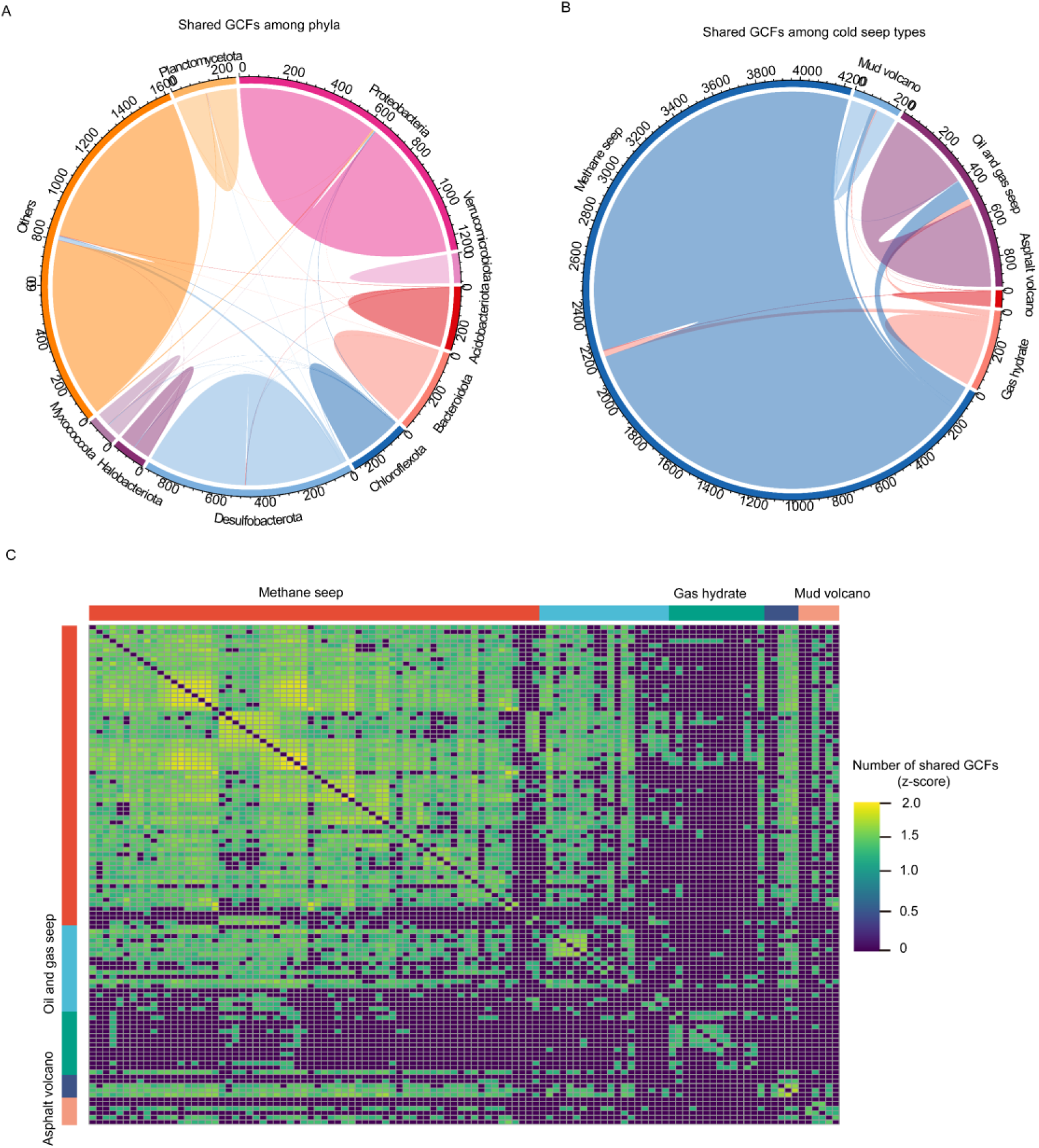
The distribution of GCFs among phyla and cold seep types. **A.** Shared GCFs within phyla (solid shapes), and with pairwise overlaps across phyla (ribbons). **B.** Shared GCFs within cold seep types. **C.** Log10-normalized pairwise heat-map of shared GCF counts among samples.

## Discussion

Although prior investigations [2,15,41–44] have focused on cold seep communities and metabolic research, a dearth of a comprehensive metagenomic-based dataset on a global scale still persists. Here, we present a specialized and fully integrated microbiome genome and gene catalogue for the global cold seep ecosystems. Compared to the previously published 1688 MAGs [15,16,19,43,44], CSMD has added 65% more genomes at the 99% ANI level, including 33 new phyla, 105 new classes, 247 new orders, 360 new families, 380 new genera, and 1094 new species (Table S11). Apart from MAGs, we also acquired un-binned contigs and integrated them with all MAGs to create a 56 Gb non-redundant contigs, a resource that has been neglected in past investigations [19,44]. This dataset is expected to be a fundament for the further exploration of evolution and gene function like glacier [10], marine [11], and human gut [12] databases.

We observed that Halobacteriota and Desulfobacterota were the dominant phyla in terms of relative abundance in cold seeps globally, which is not surprising given that they include typical ANME/SRB consortia in the cold seep [45]. Interestingly, a high abundance of Caldatribacteria was exclusively distributed in gas hydrate type, which is consistent with previous 16S-based study [5]. A recent study from the species within Caldatribacteria isolated from gas hydrates indicated environmental adaptation may link to its cell membrane structure [46]. However, whether Caldatribacteria dominates in gas hydrate type remains to be explored [5]. In addition, we found that mineral-prone systems exhibit higher alpha diversity than mud-prone systems, which is consistent with previous studies focused on viral communities [15]. This may be attributed to the longer geological history and slower fluid discharge of mineral-prone systems, providing a more stable living environment for microorganisms compared to young and fast mud-prone systems [1]. Additionally, previous studies based on 16S sequencing have shown that both sampling site and cold seep type significantly affect microbial community composition [47], and our results confirm this. Our results also indicate that sampling site has a stronger effect than cold seep type, which is not surprising considering the strong influence of environmental heterogeneity on microorganisms (small-scale spatial variation in centimeter or micrometer range may lead to dramatic changes in nutrient conditions) [48]. We discovered a rich repertoire of metabolic pathways in the cold seeps. Firstly, we found that the WL pathway is the most common carbon fixation pathway among cold seep microorganisms. Compared to the CBB and rTCA cycles, the WL pathway was lower demand for ATP, higher efficiency and faster rate [49], making it possibly the most economical choice for cold seep microorganisms. Secondly, we found that 90% of the OTUs may have the potential to degrade organic compounds based on the genes involved in carbohydrate degradation [35]. The organic compounds, including carbohydrate, were produced by AOM and settled from the upper layers of the ocean, providing a substantial nutritional status for the cold seep microbiome [1,45]. Additionally, compared to the 39% (69 out of 178 MAGs) mixotrophic ratio of microorganisms in the Challenger Deep [35], the proportion of mixotrophic microorganisms in cold seeps has increased to 61%, which may be due to the richer availability of inorganic and organic carbon sources in cold seeps [1]. Although the strict definition of mixotrophic ability is complex and usually requires experimental verification using microbial isolates, our results can be regarded as a rough, preliminary exploration.

Due to the limited availability of oxygen within a few millimeters to centimeters of the sediment surface, cold seep sediments are typically hypoxic [1]. As expected, we observed that almost all of OTUs contained at least one anaerobic metabolism pathway, indicating that anaerobic metabolism dominates in cold seeps, consistent with previous reports based on experimental and computational approaches [6,50]. Interestingly, we found that up to 39% of OTUs had the potential for facultative anaerobic respiratory capabilities. Similar results have been reported in studies with Challenger Deep [35], which may be due to the high pressure, absence of light, and low oxygen in both environments. Although more experiments are needed to verify, these results suggest that the facultative anaerobic respiratory capabilities in cold seep microbiome may have been underestimated.

The high novelty of BGCs has also been observed in marine [11] and glacier [10] environments, indicating a widespread potential for environmental microorganisms to synthesize novel natural products, which is consistent with the high novelty of environmental microbial genomes. Given that the majority of natural products currently derive from a few cultivable microbial groups [37], the high novelty of BGCs in environments such as cold seeps, which harbor a large proportion of uncultivated microorganisms, does not seem surprising. Interestingly, we found that Desulfobacterota possessed considerable biosynthetic potential, which was also observed in microorganisms from the permanently anoxic Cariaco Basin [51], suggesting that Desulfobacterota may be a lineage with unique biosynthetic encoding potential in anoxic environments. The biosynthetic potential of archaea has recently received attention [51], and we found that > 27% of archaea MAGs encoded BGCs. We anticipate that the archaea provide an even more extensive potential for novel natural products.

In summary, the CSMD provides a database and platform for archiving, analyzing, and comparing the cold seep microbiome at the genomic and genetic levels. Here, we demonstrate its unique value in exploring microbial taxonomic and functional diversity. This comprehensive work not only fills the knowledge gap in the understanding of microbial diversity and function in global cold seep ecosystems, but also provides a rich resource for natural product bioprospecting. We expect that the catalog would facilitate the research of global cold seep microbiome as more cold seep microbiome studies become available.

## Materials and methods

### Metagenomic sample collection

Overall, 113 own and public metagenomic samples were collected from different sites around the world, covering 5 different cold seep types (Figure 1A and Table S1). Among them, the SCS_HM2 dataset of 12 samples was obtained from the active Haima cold seep of South China Sea (22°07′N, 119°17′E) (Table S1) at water depth of 1100 meters during scientific cruises conducted by the research vessel “KEXUE” in 2017. Haima seeps are characterized by abundant carbonate rocks, and accompanied by a large number of living and dead bivalves [52]. Seven samples (HTR2, HTR3, HTR4, HTR5, HTR7, HTR11, and HTR12) were collected by grab sampler from the sediment surface (approximately 0–0.02 meters below seafloor (mbsf)), while three samples (HTR8, HTR9, and HTR10) were collected by Remotely Operated Vehicle (ROV) push cores from soil depths of approximately 0.02–0.2 mbsf. The remaining two samples (HTR1 and HTR6) were collected by gravity corer from soil depths of 0–1.6 mbsf. The uppermost layer impacted by seawater was discarded, and the sediment located at the core of each section was collected and stored in anaerobic biobags at –80°C for future utilization. The remaining 101 samples were downloaded from NCBI’s Sequence Read Archive (Table S1) [4–7,14,16–19,41–43].

### SCS_HM2 samples DNA extraction and metagenomic sequencing

Total DNA for SCS_HM2 sediments (∼0.5 g) were extracted using the PowerSoil DNA Isolation Kit (Catalog No. 12888-50, Qiagen, Germantown, MD) following the user’s instructions. Genomic libraries were constructed and sequenced on the Beijing Genomics Institute (BGI) MGISEQ-2000RS platform at the National Microbiology Data Center, Institute of Microbiology, Chinese Academy of Sciences (Beijing, China) with 150 bp paired-end model, followed by data processing with standard protocols.

### Metagenomic quality control and assembly

A quality control of the raw reads was performed via Trim Galore (v0.5.0) (https://www.bioinformatics.babraham.ac.uk/projects/trim_galore/). Only paired-reads with sequence lengths ≥ 100 bp were retained after adapter sequences were removed, and low quality reads were trimmed from the 3’-primer end via the Phred quality score (Q) threshold of 30. The 113 metagenomes were assembled with MEGAHIT (v1.1.3) [53] using the default k-mer parameters (--k-list 21, 29, 39, 59, 79, 99, 119, 141), retaining contigs great than 1000 bp in length. Overall mapping rate of each sample was calculated by Bowtie 2 (v2.3.5) [54] with default parameters.

### Construction of non-redundant genome and contig taxonomic annotation

The contigs derived from 113 metagenomes and public MAGs with length over 1 kb were de-replicated at 90% aligned region with 95% nucleotide identity using MMseqs2 [55] with the parameters “easy-linclust -e 0.001, -min-seq-id 0.95 and -c 0.9” [56]. Subsequently, all contigs were taxonomically annotated by CAT (v 5.2.3) [25] with default parameters based on NCBI non-redundant protein database (version 2021-01-07). The 56 Gb non-redundant contigs of the cold seep microbiome were obtained after removing eukaryotic sequences (mainly *Mytilus galloprovincialis* and *Pomacea canaliculata* that commonly accompany the Mollusca phylum in cold seep).

### Metagenome binning and genome quality control

Metagenomic assemblies were binned using MetaBAT2 (v2.12.1) [57], MaxBin2 (v2.2.7) [58] and CONCOCT [59] wrapped in MetaWRAP (v1.3.2) [60] with default parameters for each sample. In addition, VAMB (v2.0.1) [61] was also used for binning based on deep variational autoencoders. Consequently, the completeness and contamination of bins were calculated using the “lineage_wf” module of the CheckM (v1.0.12) [62]. tRNA genes were identified using the aragorn [63] and rRNA genes were identified using Barrnap (v0.9) (https://github.com/tseemann/barrnap). Finally, 4335 MAGs meeting the medium and above quality of MIMAG [24] were retained for subsequent analysis.

### Genome de-redundancy and generation of species-level OTUs

A total of 3246 previously public MAGs [15,16,19,43,44] were collected and de-redundant to 1688 genomes by dRep (v3.2.0) [64] based on genome-wide ANI percentage threshold of 99 % with parameters: -comp 50, -con 10 and -sa 0.99. Consequently, 3175 non-redundant genomes were obtained by dRep with ANI 99% combined with the previous 4335 MAGs. Finally, 1897 representative species-level OTUs were clustered using dRep based on > 30% aligned coverage, and ANI threshold of 95% (-nc 0.3, -sa 0.95) [10,13].

### MAG abundance, alpha, and beta diversity analysis

Quality-controlled reads were mapped to MAGs using minimap2-sr [65] with default parameters. The abundance of MAGs was calculated using CoverM (version 0.6.0) (https://github.com/wwood/CoverM) with parameters: --min-read-aligned-percent 0.75, - -min-read-percent-identity 0.95, --proper-pairs-only, --methods tpm. Transcripts Per Million (TPM) was used to eliminate the effects of sample sequencing depth and genome length [10,66]. In addition, PhyloFlash (v3.4.1) [67] was used for extracting 16S miTags from clean metagenomic data using parameters “-almosteverything”, and classifying via SILVA database (v138.1) [68]. Subsequently, the “rarecurve” function in the vegan package (https://github.com/vegandevs/vegan/) of R was used to assess sample sequencing saturation to remove samples with low sequencing depth. Taxonomic structure plot, alpha, and beta diversity analyses were performed using the R package EasyMicroPlot [69]. Mann-Whitney test was used for two groups of Shannon as well as Simpson indices, and one-way analysis of variance (ANOVA) and Tukey HSD post-hoc tests were used among groups [70]. For beta diversity analysis, Bray-Curtis distances were measured, and PERMANOVA analysis was used to test for statistical significance among different independent variables with the default settings (999 permutations).

### Comparison MAGs to genomes of public databases

The species-level representative OTUs were compared to 103,722 publicly available reference genomes, including 968 genomes from the TGG [10], 957 MAGs from the TaraOcean [20], 8304 MAGs from the OMD [11], 45,599 MAGs from the GEM [13], and 47,894 MAGs from the GTDB [29]. Each reference dataset was compared with 1897 OTUs using dRep. A cold seep OTUs was designated as novel species which exhibit an ANI less than 95% with other reference genomes [10].

### Metabolic pathway analysis of MAGs

Genes were predicted for MAGs using Prokka (v1.14.6) [71] with single genome model. KEGG pathway was then annotated by eggNOG-mapper (v2.1.6) [72] based on eggNOG Orthologous Groups database (version 5.0) [73]. To elucidate an overview of the specific metabolic modules of each MAGs, the key enzymes of the metabolic pathway are summarized and visualized follow the method of Chen et al [35]. The module completeness of a given metabolic pathway can be quantified as the percentage of identified key marker genes present in the corresponding pathway. For example, a module completeness value of 50 indicates that the MAG contains 50% of the marker genes in the complete pathway [35].

### Taxonomic annotation and phylogenetic tree inference

The taxonomic annotation of the 3175 MAGs were performed using the Genome Taxonomy Database Toolkit (GTDB-Tk, v0.3.2) [30] with the GTDB database release R06-R202 [47]. MAGs were classified at the species level if the ANI to the closest GTDB-Tk representative genome was ≥ 95% and the aligned coverage was ≥ 60%. Finally, the phylogenetic tree was inferred by IQ-TREE (v2.2.0-beta) [74] with parameters: -B 1000, -m LG + G, -wbtl, based on the concatenated multiple sequence alignments of 122 archaeal, or 120 bacterial universal marker genes generated by GTDB-Tk after trimming sequence gaps via trimAl (v1.4.rev15) [75]. iTOL [76] was used to visualized phylogenetic trees.

### Gene function annotation

The prediction of open reading frames (ORFs) in metagenomic assemblies was carried out using Prodigal [49]. The resulting ORFs were then dereplicated by clustering at 80% aligned region with 95% nucleotide identity, employing MMseqs2 [50] with the parameters: easy-linclust -e 0.001, --min-seq-id 0.95, -c 0.80 [10]. The gene rarefaction analysis was performed using an in-house python script, based on the gene cluster result of MMseqs2 with identity thresholds of 95%, 75%, and 50% (easy-linclust -e 0.001 -c 0.80) [10]. The analysis was repeated 100 times with a 5% sampling step. Further, the function of the non-redundant gene catalog was annotated to the Swiss-Prot [27], Uniref 50 [28], and NR (ftp://ftp.ncbi.nlm.nih.gov/blast/db) databases via MMseqs2 [55] with parameters: easy-search -e 0.01, --min-seq-id 0.3, --cov-mode 2 -c 0.8.

### The secondary metabolite BGC analysis

The secondary metabolite BGC was predicted for contigs ≥ 3 kb in length using AntiSMASH (v5.1) [36] with default settings. Subsequently, the BGCs were categorized into GCFs and labeled with seven categories: ‘PKSI’, ‘PKS-NPR_Hybrids’, ‘PKSothers’, ‘NRPS’, ‘RiPPs’, ‘Terpene’, and ‘Others’, based on the results of the BiG-SCAPE [21] with default parameters.

### Novelty of GCFs

The novelty of GCFs was estimated based on the result of BLASTN (BLAST 2.2.28+) [40,77] to databases of experimentally validated MIBIG 2.0 [37] and the latest computationally predicted OMD [11]. For the representative BGC of GCFs, we selected the sequence with maximum query coverage and identity to the respective database as the best hit. A GCF was deemed novel if the best hit to the reference was below 80% query coverage and 75% identity following the threshold of GEM [13].

## Data availability

Raw reads of 12 samples in this study are deposited in the NCBI Sequence Read Archive (SRA) under accession PRJNA916811, the National Microbiology Data Center (NMDC, https://nmdc.cn/) under accession number NMDC10018281, and also available from Genome Sequence Archive [78] in National Genomics Data Center (NGDC), China National Center for Bioinformation/Beijing Institute of Genomics (BIG), Chinese Academy of Sciences under accession CRA010074 that are publicly accessible at https://ngdc.cncb.ac.cn/gsa. The genome sequences of 3175 MAGs and 54 Gb non-redundant contigs of CSMD are deposited in Genome Warehouse (GWH) [79] in NGDC under accession PRJCA015385, and also available from https://doi.org/10.6084/m9.figshare.21731330.

## CRediT authorship contribution statement

**Tao Yu:** Conceptualization, Methodology, Investigation, Formal analysis, Writing - original draft. **Yingfeng Luo:** Conceptualization, Writing - review & editing. **Xinyu Tan:** Methodology. **Dahe Zhao:** Methodology. **Xiaochun Bi:** Methodology. **Chenji Li:** Methodology. **Yanning Zheng:** Methodology, Writing - review & editing. **Hua Xiang:** Conceptualization, Writing - review & editing. **Songnian Hu:** Conceptualization, Writing - review & editing, Supervision. All authors have read and approved the final manuscript.

## Acknowledgments

We thank the Senior User Project of RV KEXUE (No. KEXUE2019GZ05), and thank the Center for Ocean Mega-Science, Chinese Academy of Sciences. We acknowledge the data and samples collection by RV KEXUE. We also thank the Second Tibetan Plateau Scientific Expedition and Research Program (2021QZKK0100 (Y. Luo and T. Yu) and the National Key Research and Development Plans 2022YFF1002801 (Y. Luo).

## Supplementary materials

**Figure S1.**
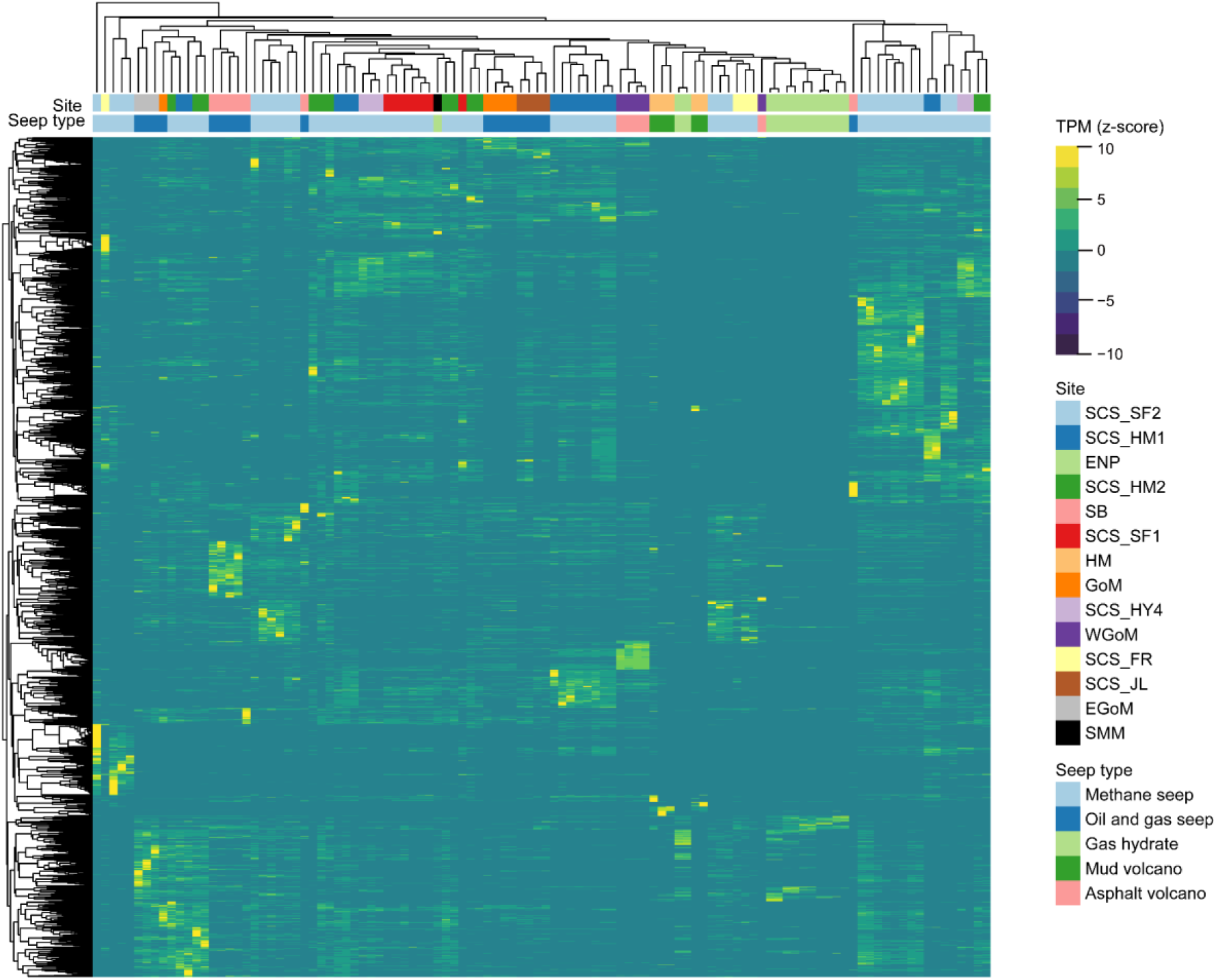
Heat-map presentation of relative abundance of CSMD 1897 OTUs among samples. The abundance of 1897 OTUs in 113 samples normalized by z-sore, and bi-directional clustering. Cold seep types are shown as column annotation, and each OTU is shown as row annotation.

**Figure S2.**
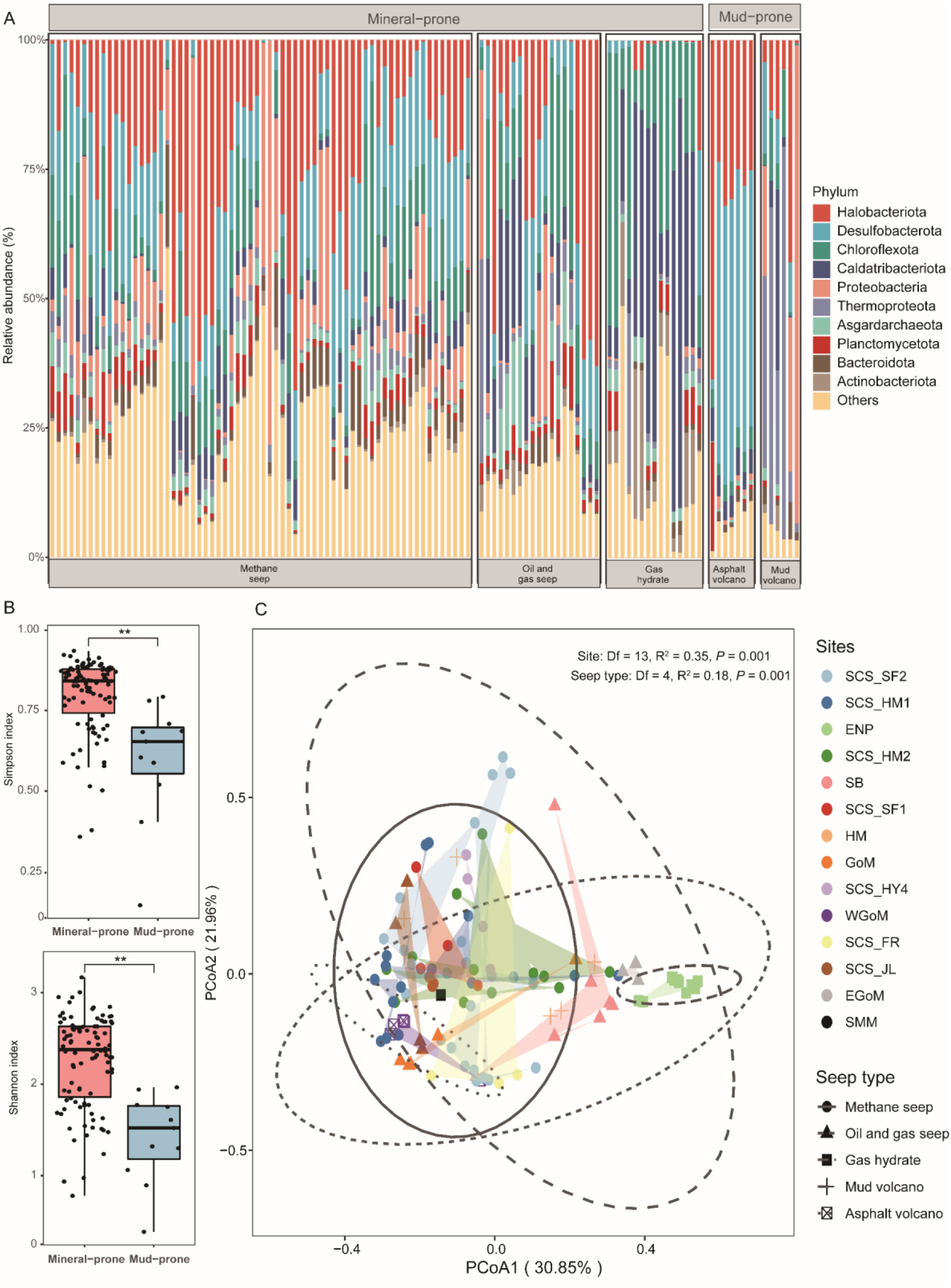
Cold seep microbiome composition barplot, alpha and beta diversity based on MAGs abundance at phylum level. **A.** Taxonomic relative abundance barplot of cold seep microbiome. **B.** Simpson and Shannon diversity between mineral-prone system and mud-prone system. Statistics by Mann−Whitney test, ** indicate significance at *P* < 0.01. **C.** Beta diversity difference among cold seep sites and types based on Bray-Curtis dissimilarities. Ellipses represent 95% confidence contours of samples grouped by cold seep type. PERMANOVA analysis was used to test for statistical significance for the main effects of cold seep sites and types.

**Figure S3.**
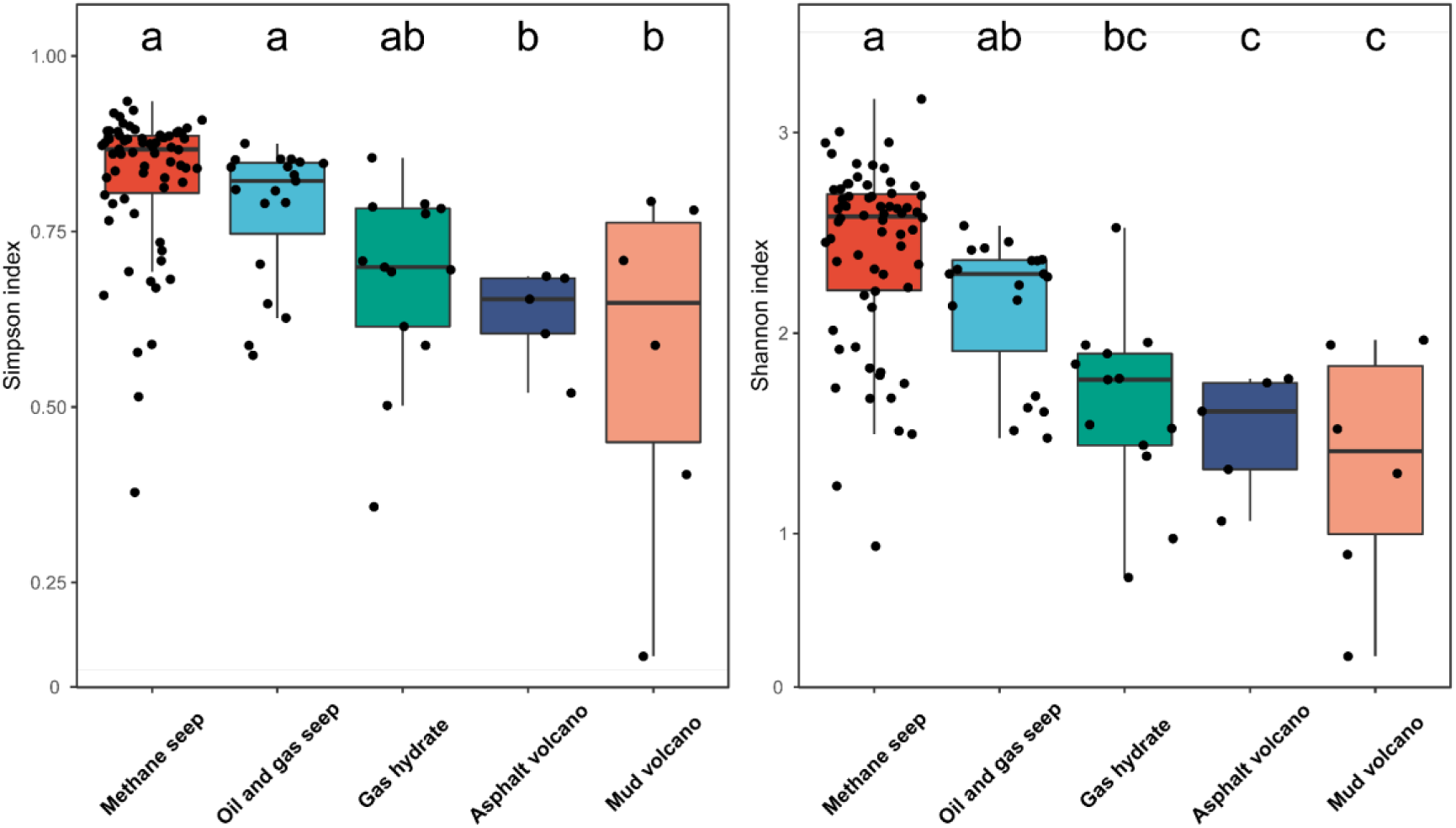
Simpson and Shannon diversity among cold seep types. Different letters indicate the values that differ significantly among cold seep types at *P* < 0.05 (one-way analysis of variance (ANOVA), and Tukey HSD post-hoc tests).

**Figure S4.**
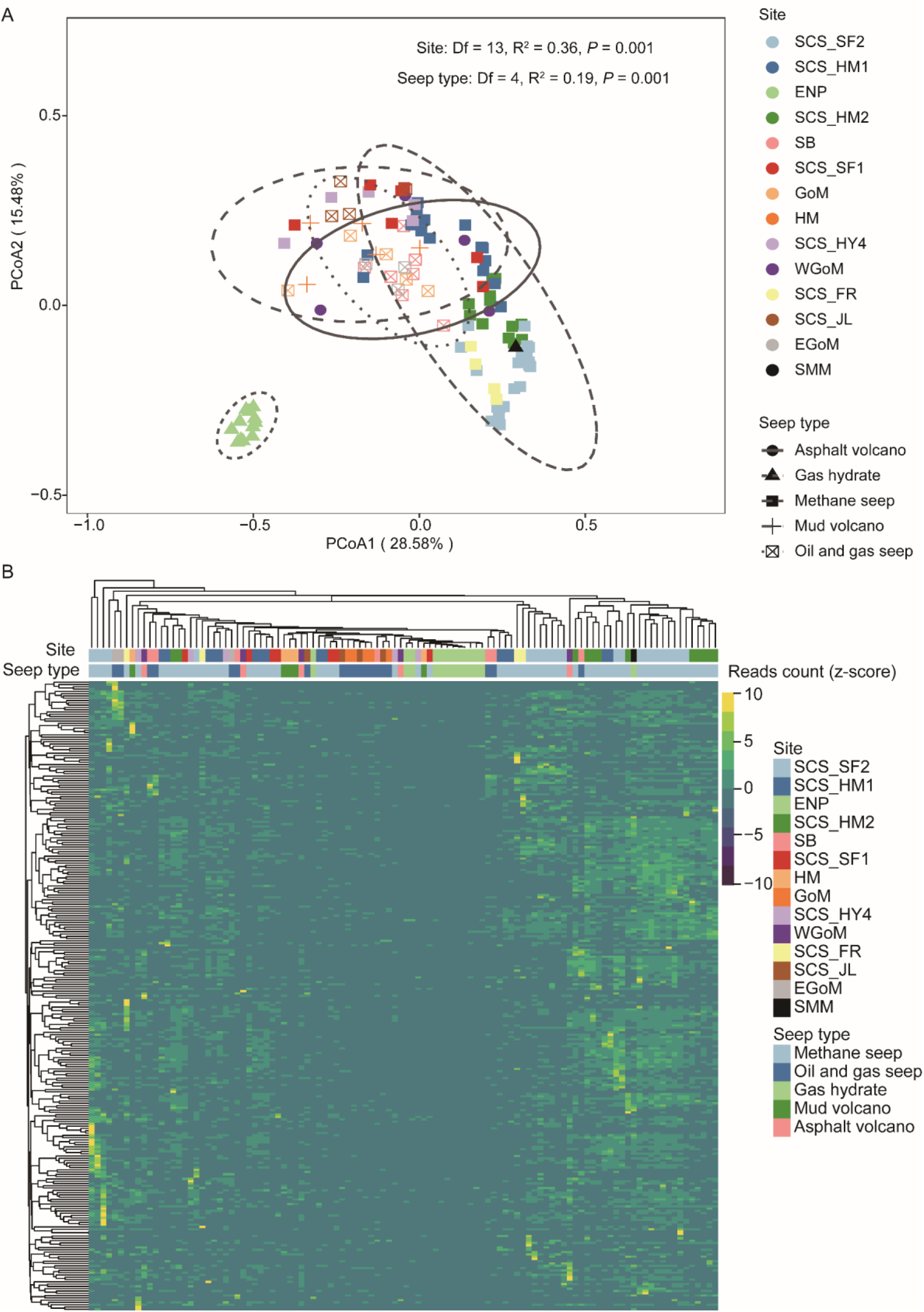
Cold seep microbiome beta diversity and heat-map based on 16S abundance at phylum level. **A**. Beta diversity difference among cold seep sites and types based on Bray-Curtis dissimilarities. Ellipses represent 95% confidence contours of samples grouped by cold seep type. PERMANOVA analysis was used to perform statistical significance for the main effects of cold seep sites and types. **B**. The heat-map of 16S abundance with normalized by z-score, and bi-directional clustering. Cold seep types are shown as column annotation, and each OTU is shown as row annotation.

**Figure S5.**
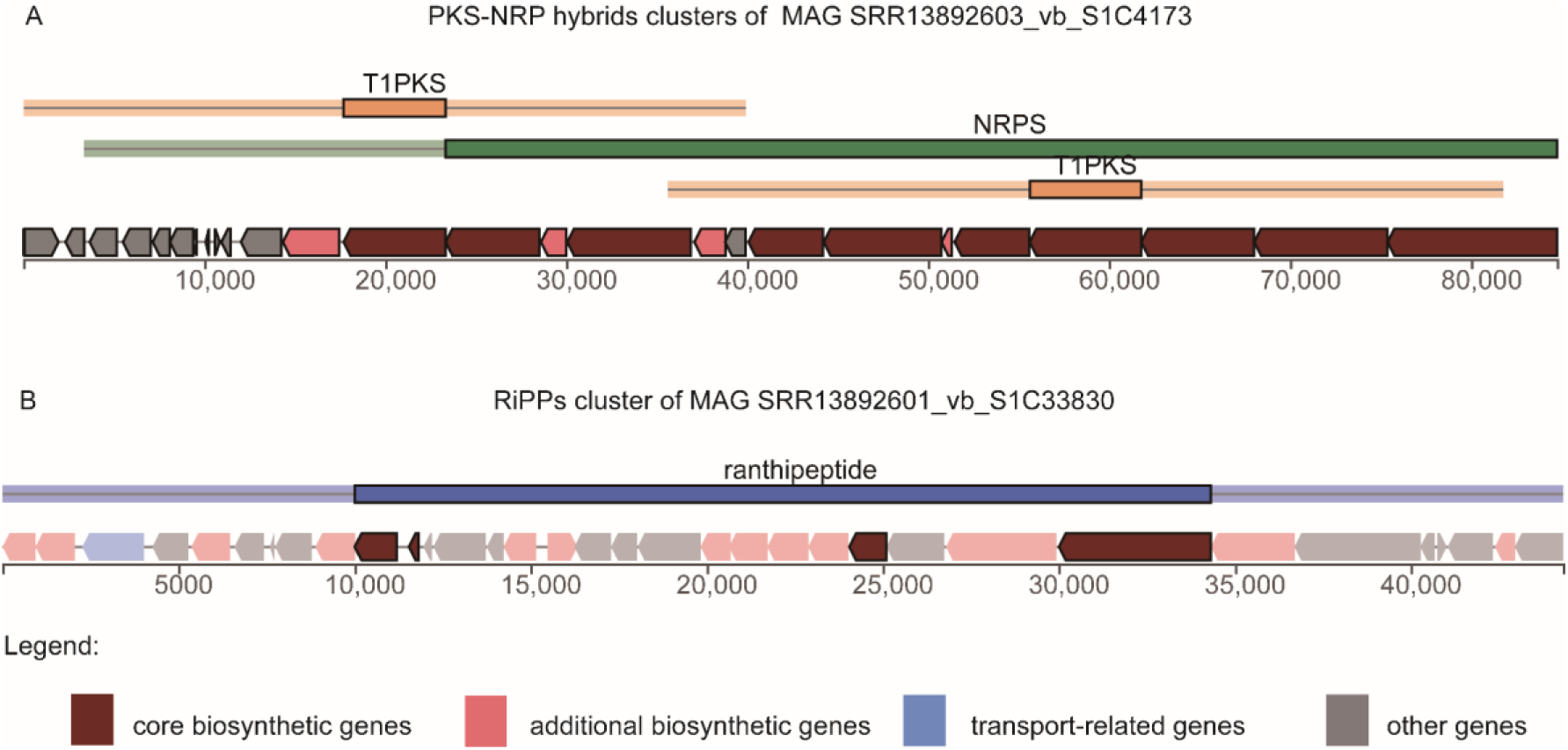
The genomic structures of two identified BGCs. BGCs predicted from a novel genus (SRR13892603_vb_S1C4173) and a species (SRR13892601_vb_S1C33830) encode a PKS-NRP hybrid and a RiPP gene cluster, respectively. **A.** The PKS-NRP hybrid cluster of SRR13892603_vb_S1C4173 comprises 10 core modules and spans 84,733 bp. **B.** The RiPP cluster of SRR13892601_vb_S1C33830 comprises 4 core modules and spans 44,319 bp.

**Table S1 The global cold seep metagenome and assembly statistics**

**Table S2 The genomic characteristics of 3175 MAGs**

**Table S3 The read mapping rate of self-assembly and 56 Gb non-redundant contigs**

**Table S4 The novelty comparison of CSMD OTUs with public genomes**

**Table S5 The shared OTU members among cold seep types**

**Table S6 The OTU abundance among samples**

**Table S7 Taxonomic expansion of CSMD compared with GTDB-RS202 at the phylum level**

**Table S8 Reads count of 16S (miTags) among samples**

**Table S9 The metabolic profile of 1897 OTUs**

**Table S10 A summary of 17,968 BGCs**

**Table S11 The BGC class count of MAGs**

**Table S12 Taxonomic expansion of CSMD compared to public MAGs of cold seeps**

